## Supplementary material for "Two stages of bandwidth scaling drives efficient neural coding of natural sounds": Supplmental Text 1

**Supporting Information**

Here we derive the relationship between STRF domain parameters and the corresponding modulation domain parameters that are necessary to produce scaling. The scaling characteristics of the model STRFs are achieved by requiring that the quality factor of the modulation filters is ~1, thus consistent with auditory midbrain measurements [1] and human perceptual studies [2,3].

The modulation transfer function (MTF) represents a neuron's sensitivity in the modulation domain. It is the two-dimensional Fourier transform of the neuron's STRF[4]

$$MTF(f_m, \Omega) = \iint STRF(t, x)\, \mathrm{e}^{-j2\pi(\Omega x + f_m t)} dt dx$$

where Ω is the spectral modulation frequency (also widely referred as ripple density) which represents the number of spectral oscillations per unit octave (cycles/octave) and $f_m$ is the temporal modulation frequency (Hz). For the model Gabor STRFs of Eqn. 6, the MTF magnitude is a separable function

$$|MTF(f_m, \Omega)| = |MTF_S(\Omega)| \cdot |MTF_t(f_m)|$$

where

$$|MTF_S(\Omega)| = e^{-\left[\frac{\pi bw(\Omega - \Omega_o)}{2}\right]^2}$$

is the spectral MTF magnitude and

$$|MTF_t(f_m)| = \frac{1/\tau^2}{4\pi^2(f_m - f_{m0})^2 + 1/\tau^2}$$

is the temporal MTF magnitude.

The peak of the MTF is determined by $f_{m0}$ and $\Omega_0$ while the bandwidth is measured at 3dB (50% power) for both temporal and spectral dimensions. Since these two dimensions are separable, the modulation bandwidths can be measured separately for each. Considering a ½ power criteria

$$|MTF_S(\Omega)|^2 = \left(e^{-\left[\frac{\pi bw(\Omega - \Omega_0)}{2}\right]^2}\right)^2 = 0.5$$

the 3-dB cutoff frequencies and spectral modulation bandwidths are

$$\Omega_{3dB} = \Omega_o \pm \frac{2\sqrt{\frac{1}{2}ln(2)}}{\pi \cdot bw} = \Omega_o \pm \frac{\sqrt{2ln(2)}}{\pi \cdot bw}$$

$$\mathrm{BW}_\Omega = \frac{2 \cdot \sqrt{2 \cdot ln(2)}}{\pi \cdot bw}.$$

Similarly, for the temporal dimension solving the ½ power criteria

$$|MTF_t(f_m)|^2 = \left(\frac{1/\tau^2}{4\pi^2(f_m - f_{m0})^2 + 1/\tau^2}\right)^2 = 0.5$$

yields the 3-dB cutoff frequency and temporal modulation bandwidth

$$f_{m3dB} = f_{m0} \pm \frac{1}{2\pi \cdot \tau}\sqrt{\sqrt{2} - 1}$$

$$\mathrm{BW}_{fm} = \frac{\sqrt{\sqrt{2}-1}}{\pi} \cdot \frac{1}{\tau}.$$

Note that under these constraints, temporal ($f_{m0}$) and spectral ($\Omega_o$) modulation bandwidths are independent of the best temporal and spectral modulation frequencies.

Next, we consider the fact that modulation filters in the auditory midbrain have a quality factor of ~1 for both spectral and temporal modulations. Thus, under this constraint ($Q_\Omega = \Omega_0/BW_\Omega = 1$ and $Q_{f_m} = f_{m0}/BW_{f_m} = 1$) which requires that the modulation bandwidths are equal to the best modulation frequency ($BW_{f_m} = f_{m0}$ and $BW_\Omega = \Omega_0$). Under these constraints, $\tau$ and $bw$ are now inversely related to $\Omega_0$ and $f_{m0}$, respectively. For the spectral domain

$$\mathrm{BW}_\Omega = \Omega_0 = \frac{2 \cdot \sqrt{2 \cdot ln(2)}}{\pi \cdot bw}$$

so that

$$bw = 2\frac{\sqrt{2 \cdot ln(2)}}{\pi \cdot \Omega_0}.$$

Similarly, for the temporal domain

$$\mathrm{BW}_{fm} = f_{m0} = \frac{\sqrt{\sqrt{2}-1}}{\pi} \cdot \frac{1}{\tau}$$

and thus

$$\tau = \frac{\sqrt{\sqrt{2}-1}}{\pi} \cdot \frac{1}{f_{m0}}.$$

This proof thus demonstrates that the STRF and modulation domain parameters are inversely related, so long as the modulation and STRF-domain functions exhibit scaling similar to auditory midbrain.

Supporting Figures

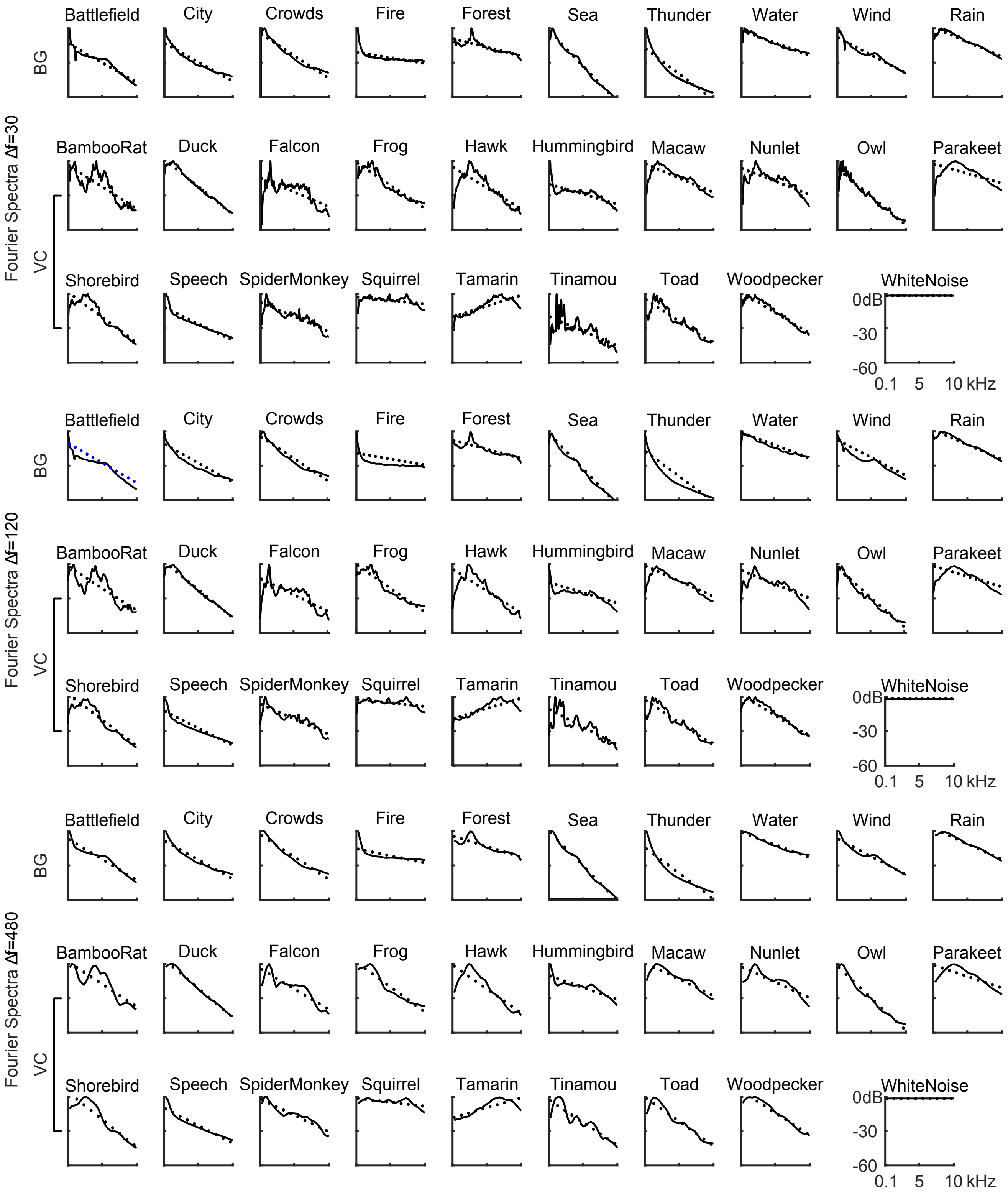


Figure S1. Fourier power spectra for natural sounds with different resolutions. Power spectra for all sound categories are analyzed using the Fourier-based model with resolutions: 30, 120 and 480Hz.

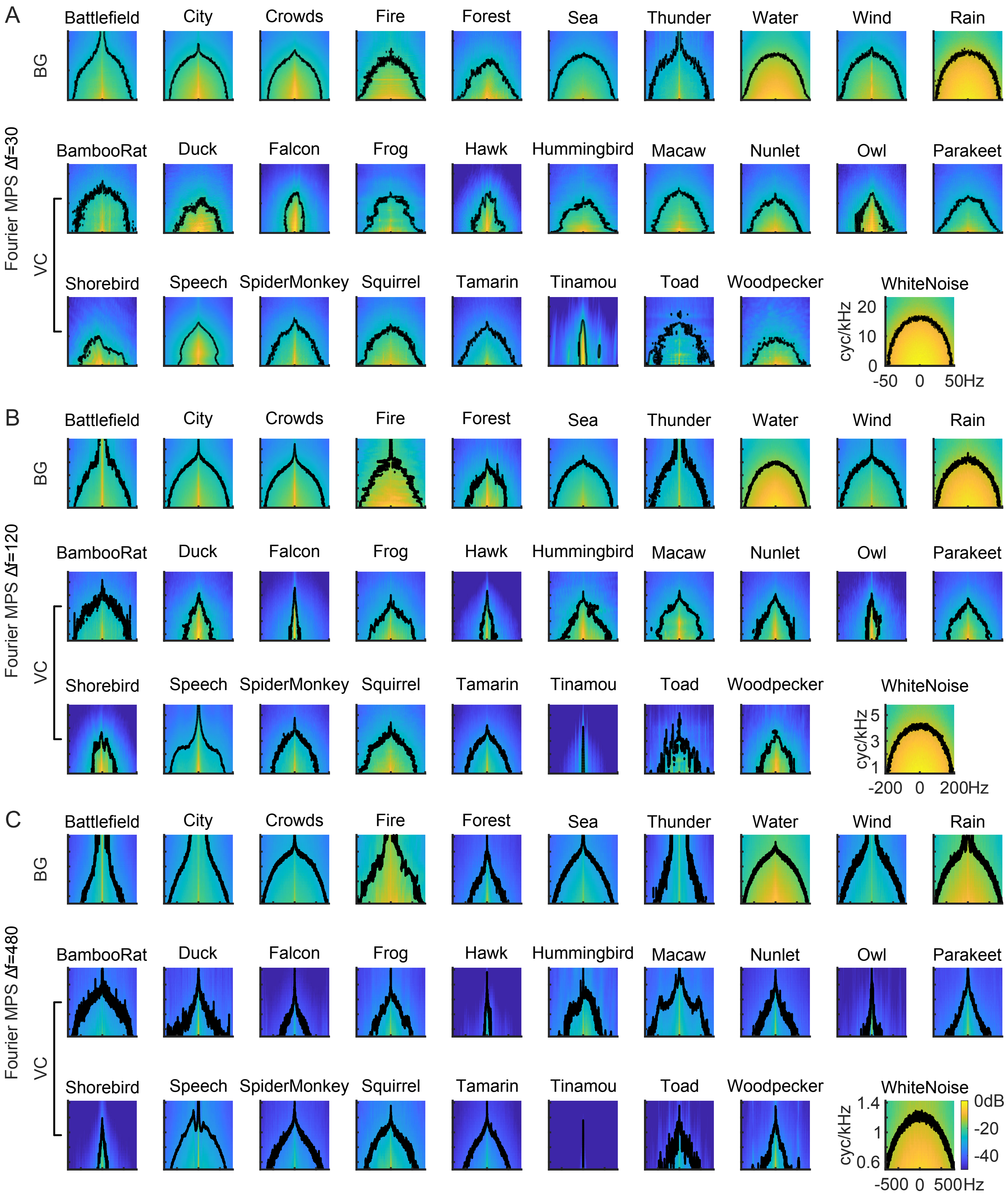


Figure S2. Fourier modulation power spectra of natural sounds with different resolutions. Modulation power spectra for all sound categories are analyzed with the Fourier model with resolutions: 30, 120 and 480Hz.

AUDIO COMPILATIONS

Audio clips were obtained from commercially available CD compilations and websites, as noted below. Details of each individual sounds used (source, track number, duration etc.) are provided in S1_Table.


1. Sony Pictures Sound Effects Series: Sony Corporation; 2003.
2. Atmospheres & Environments Sound Effects: Sound Ideas.
3. Emmons LH, Whitney BM, Ross DL. Sounds of Neotropical Rainforest Mammals: An Audio Field Guide. Ithaca, NY: Macaulay Library of Natural Sounds, Cornell Laboratory of Ornithology; 1997.
4. Sounds of Nature & The Great Outdoors: Madacy Records; 1994.
5. Stokes D, Stokes L. Stokes Field Guide to Bird Songs: Western Region2010.
6. Schulenberg TS. Voices of Amazonian Birds, Vol. 1: Tinamous Through Barbets: Cornell Laboratory Of Ornithology; 2000.
7. Davidson C. Composer, Frog and Toad Calls of the Rocky Mountains: Vanishing Voices.: Cornell Laboratory Of Ornithology.; 1996.
8. LibriVox. Available from: https://librivox.org.
9. Whitney BM, Parker TA, Budney GF, Munn CA, Bradbury JW. Voices of New World Parrots: Cornell Laboratory Of Ornithology; 2002.
10. Lipitz M. A walk in the forest: The Music Company; 1993.
11. Storm J. Earthtunes Storms in the Smokies: The Library of Natural Sounds - Cornell Library of Ornithology; 1994.
12. Sound Ideas General Catalog: Sound Ideas. Available from: https://www.sound-ideas.com.
